# A role for CREB5 in skin wound healing

**DOI:** 10.64898/2026.08.21.746254

**Authors:** Chen Han, Heidi Yuan, Trevor R. Leonardo, Kimberly Glass, Lin Chen, Luisa A. DiPietro

**Affiliations:** Center for Wound Healing and Tissue Regeneration, University of Illinois Chicago, Chicago, IL, USA; Department of Integrative and Translation Physiology, University of Illinois Chicago, Chicago, IL, USA; Department of Microbiology and Immunology, University of Illinois Chicago, Chicago, IL, USA; Channing Division of Network Medicine, Brigham and Women’s Hospital, Boston, MA, USA and Harvard Medical School, Boston, MA, USA

## Abstract

Compared with skin wounds, oral mucosal wounds heal more quickly, with minimal scarring, faster re-epithelialization, and reduced inflammation. One differentiating factor may be the differential transcription factor-associated gene networks involved in tissue regeneration. One such transcription factor, BATF3, was recently shown by us to promote wound-healing responses in vitro and in vivo. Our prior analyses also suggest that CREB5 is a differentially regulated transcription factor in oral wounds and may be involved in early wound-healing gene expression programs. CREB5 expression was induced in immortalized skin keratinocytes (HaCaT) to examine its effect on in vitro wound healing relative to immortalized gingival keratinocytes (TIGK). CREB5 overexpression let to differential expression of predicted downstream genes and improved skin keratinocyte migration in vitro. This work suggests that examining transcription factors and gene networks that regulate wound-healing responses in the oral mucosa may lead to the discovery of novel targets to improve skin wound healing.

## Introduction

Oral mucosal and skin wounds exhibit site-specific differences in their transcriptomic responses to injury [1–7]. Several transcription factors have already been identified to be differentially expressed in oral wounds relative to skin, including BATF3, PITX1, and SOX2 [1, 6, 7]. We recently demonstrated that BATF3 accelerates wound-healing responses *in vitro* and *in vivo* [7]. Another transcription factor predicted to contribute to enhanced oral wound regeneration is CREB5 [6–8]. We hypothesize that CREB5, similar to BATF3, may enhance skin keratinocyte wound healing when induced.

## Methods & Materials

### i. Cell culture

Spontaneously immortalized skin keratinocytes (HaCaT) (AddexBio, San Diego, CA, USA, Catalog #T0020001) were propagated in Dulbecco’s modified Eagle’s medium (DMEM) (4.5 g/L glucose, sodium pyruvate, L-glutamine) (Mediatech, Manassas, VA, USA), supplemented with 10% FBS (GeminiBio, Sacramento, CA, USA) and penicillin-streptomycin (0.1%) (Life Technologies, Carlsbad, CA, USA). Cells were incubated at 37 °C in a humidified atmosphere containing 5% CO2 and used for *in vitro* assays when they reached 70-90% confluence.

### ii. DNA plasmid constructs

Green fluorescent protein (GFP) was used as the reporter gene for this study. Transfected plasmids include the pLenti-CMV-Blank-CBH-GFP-2A-Puro Control Vector (Applied Biological Materials [ABM], Richmond, BC, CAN, catalog no. LV590) and pLenti-III-CMV-hCREB5-GFP-2A-Puro (ABM, catalog no. 168150610395). Plasmids were transfected into DH5α competent E. coli (Zymo Research, CA, USA), which were then grown overnight in LB medium (MP Biomedicals, Santa Ana, CA, USA, catalog no. 113002121-CF) supplemented with 50 μg/mL kanamycin. The plasmid was purified for transfection with a ZymoPURE II Plasmid Midiprep Kit (Zymo Research).

### iii. Polyethylenimine plasmid DNA transfection

HaCaT were seeded into 6-well plates and grown until 70-80% confluent. Twenty-four hours before transfection, the HaCaT growth medium was replaced with antibiotic-free medium. A polyethylenimine (PEI) plasmid DNA (pDNA) transfection mix was prepared by combining 1 µg/mL of linear molecular weight 250,000 PEI, (Polyscience, Niles, IL, USA) with plasmid (pDNA) at a mass ratio of 1:2 µg of PEI:pDNA. PEI and pDNA were added to separate 250 µL aliquots of Opti-MEM (ThermoFisher Scientific, Waltham, MA, USA) and incubated for 20 minutes at room temperature. The PEI Opti-MEM solution was added dropwise to the pDNA Opti-MEM solution and incubated at room temperature for 30 minutes. The PEI-pDNA nanoparticle transfection mixture was then added dropwise to HaCaT and incubated overnight.

### iv. Isolation and selection of positively transfected HaCaT

Following plasmid transfection, HaCaT cultures were washed with PBS, trypsinized, and centrifuged at 300 x g for 5 minutes. The cell pellet was resuspended in DMEM (Mediatech) with 20% FBS (GeminiBio) containing 7-amino-actinomycin D (BD Biosciences, Franklin Lakes, NJ, USA) to exclude dead cells, and subjected to cell sorting analysis using MoFlo Astrios Cell Sorter (Beckman Coulter, Inc., Indianapolis, IN, USA) to obtain positively transfected HaCaT. Two HaCaT cell lines were obtained: GFP-expressing Empty Vector Control HaCaT (EV Ctrl) and GFP-expressing CREB5 Overexpressing HaCaT (CREB5 OvExp).

EV Ctrl HaCaT and CREB5 OvExp HaCaT were incubated in normal growth media for 24 hours before antibiotic selection with 2.5 µg/mL puromycin for 48 hours. Following selection, cells were washed with PBS and cultured in standard growth medium supplemented with 0.25 µg/mL puromycin. EV Ctrl and CREB5 OvExp HaCaT were used for *in vitro* assays once they reached 80-90% confluency.

### v. Real-Time PCR Analysis

To confirm overexpression of *CREB5*, real-time PCR (RT-PCR) analysis was performed. EV Ctrl and CREB5 OvExp HaCaT were grown in a 12-well plate. Once confluent, total RNA was extracted using TRIzol (Invitrogen, Waltham, MA, USA). One µg of total RNA was treated with DNase (ThermoFisher Scientific) and converted to cDNA using a High-Capacity cDNA Reverse Transcription Kit (Invitrogen). Relative expression of *CREB5* and target genes was determined by semi-quantitative PCR on a StepOnePlus RealTime PCR System (Applied Biosystems, Waltham, MA) using Power SYBR Green PCR Master Mix (Roche, Basel, SUI). Relative gene expression was determined with the 2−ΔΔCT method [9]. Beta-actin (*ACTB*) was used as a reference gene, and EV Ctrl was used as the control. Supplementary Table 1 shows the genes and primers used for our RT-PCR analyses. *CREB5* target genes were identified in a similar fashion as described for BATF3 in our previous study [7]

### vi. Analysis of in vitro cell migration and proliferation

To determine if *CREB5* overexpression enhances HaCaT phenotypes related to wound healing, we performed *in vitro* functional assays using EV Ctrl and CREB5 OvExp HaCaT. All experiments were repeated at least 2 times. For the migration assay, we used an *in vitro* wounding model in which cells were grown to confluence in a 12-well plate and then treated with mitomycin-C (1 μg/ml) (Sigma-Aldrich, St. Louis, MO, USA) for 1 hour to inhibit proliferation. *In vitro* wounding was performed by scraping the plate horizontally and vertically with a 200 µL pipette tip, creating 1×1 cross-scratches in each well. After wounding, cells were washed with PBS, and fresh media was added.

Open areas were photographed at 0, 24, and 48 hours after wounding and measured using ImageJ [10]. Migration rate is expressed as a percentage of the original uncovered area at each time point. To compare EV Ctrl and CREB5 OvExp HaCaT proliferation, we seeded 5000 cells into each well of a 96-well plate. Proliferation was assessed at 48 and 96 hours post-seeding using an MTS Cell Proliferation Assay Kit (Abcam, Waltham, MA, USA, Catalog # ab197010). Optical density (OD) values were measured at 490 nm with a spectrophotometer (Molecular Devices, San Jose, CA, USA).

## Results

### CREB5 improves HaCaT wound healing phenotype

*CREB5* is a transcription factor predicted to be a top regulator in gene networks of early palate wound response (Figure 1A) [6, 7]. A positive correlation indicates that increased *CREB5* targeting is associated with increased target gene expression.

**Figure 1.**
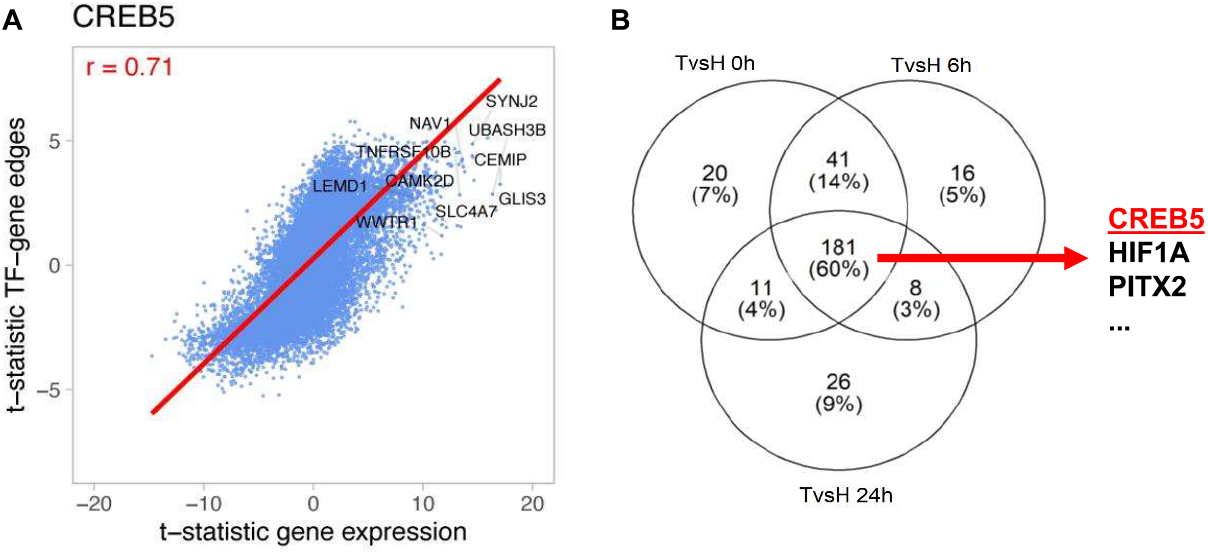
CREB5 is a significant transcriptional activator in palatal versus skin wounds. A) Scatter plot (blue) and regression line (red) of the differential targeting of CREB5 with its putative target genes versus the differential expression of those genes in the 6 h vs. 0 h LIMMA network and differential expression comparisons in palate. Each point represents a gene that is a putative CREB5 target based on the motif prior. The x-axis represents the t-statistics of the differential expression when comparing 6 h post-injury vs. unwounded palate samples. The y-axis represents the t-statistics of the differential targeting (edge weight) when comparing 6 h post-injury vs. unwounded palate gene regulatory networks. A positive t-statistic on the x- or y-axis represents increased gene expression or increased CREB5 targeting of that gene at 6 h compared to unwounded tissue, respectively. A negative t-statistic on the x- or y-axis represents decreased gene expression or CREB5 targeting of that gene at 6 h compared to unwounded tissue, respectively. A Pearson’s correlation coefficient of r= 0.71, labeled in red, corresponds to CREB5’s regulatory activity in palate at 6 h post-injury. The target genes with the most significant differential targeting that are also significantly overexpressed (FDR<0.05) at 6 h after injury are labeled in black. B) Venn diagram highlighting the transcription factors that are always up-regulated in TIGK vs HaCaT (TvsH) at 0 h, 6 h, and 24 h post-injury.

Interestingly, *CREB5*, along with other transcription factors known to regulate migration and re-epithelialization, was consistently elevated in TIGK relative to HaCaT throughout healing (Figure 1B) [1, 6, 11, 12]. To assess whether *CREB5* expression augments the keratinocyte wound-healing phenotype, we generated HaCaT cell lines overexpressing CREB5 (Figure 2A) and evaluated the expression of the top genes predicted to be targeted by *CREB5* (Figure 1A). Several predicted CREB5 target genes exhibited significant changes in expression in CREB5-overexpressing HaCaT cells compared with control cells (Figure 2B). Specifically, *CAMK2D and TNFRSF10B* were upregulated, while *SYNJ2* and *UBASB3B* were downregulated (Figure 2B) [7]. As compared to control, *CREB5-*overexpressing HaCaT did not exhibit significantly different rates of proliferation but did exhibit more rapid *in vitro* wound closure (Figure 2C-E). Altogether, our results suggest that *CREB5* overexpression regulates HaCaT gene networks and, subsequently, enhances the HaCaT wound-healing phenotype *in vitro*.

**Figure 2.**
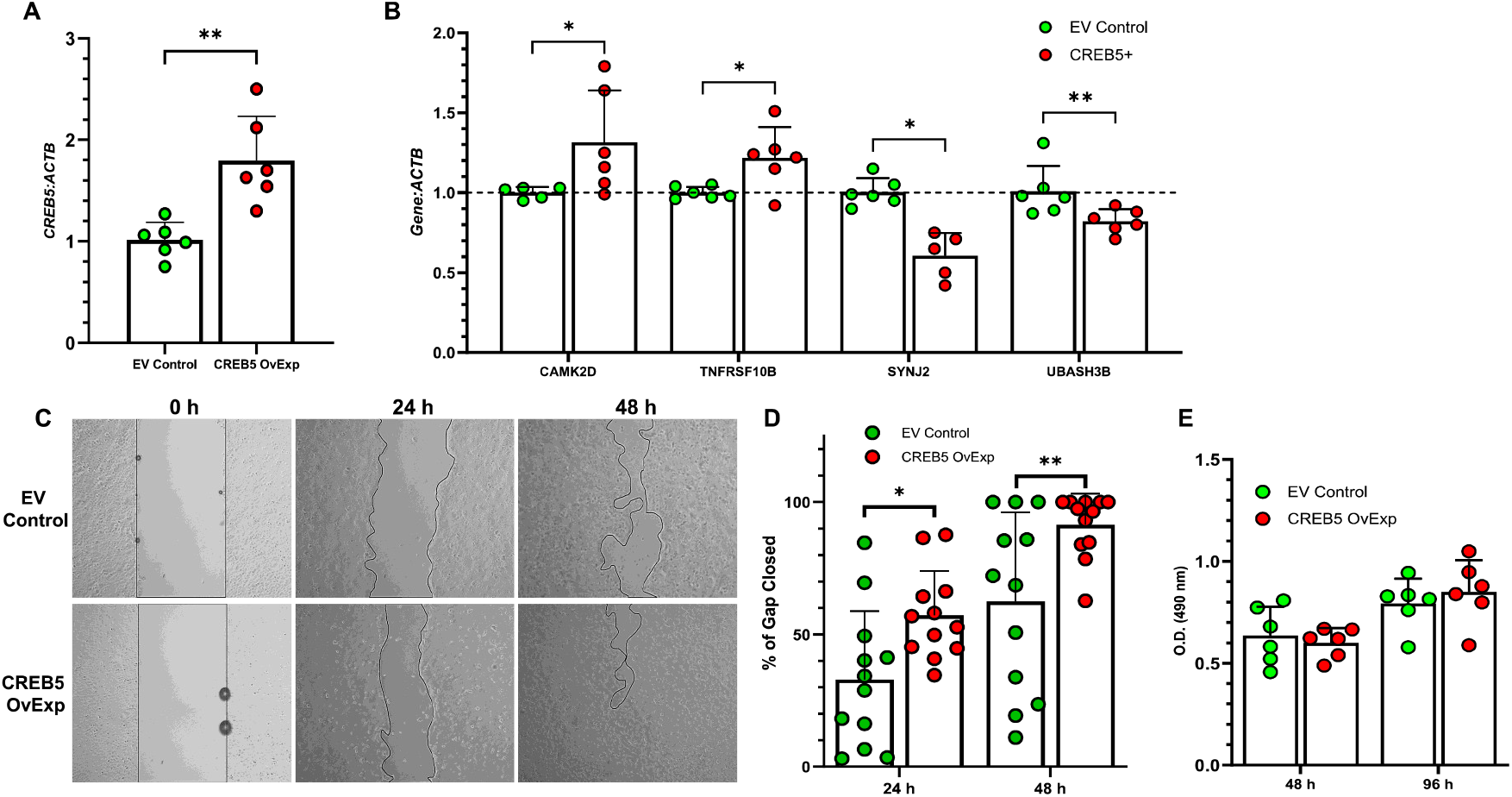
CREB5 improves HaCaT migration *in vitro*. A) Gene expression levels of *CREB5* in the empty vector control (EV Control) and *CREB5* overexpression vector (CREB5 OvExp) HaCaT cell lines. N=6, with each dot representing a biological replicate that consists of 3 technical replicates. B) Gene expression levels of putative CREB5 targets normalized to *ACTB* in EV Control and CREB5 OvExp. N=5-6, with each dot representing a biological replicate that consists of 3 technical replicates. C) Representative photos of *in vitro* vertical EV Control and CREB5 OvExp wounds closing after one x one cross-scratching. A black line outlines areas not covered by cells. D) Rate of cell migration for EV Control and CREB5 OvExp is expressed as a percentage of the original uncovered scratch area (N=12). E) Proliferation of EV Control and CREB5 OvExp at 48 and 96 h post-seeding was assessed by MTS assay. N=6, with each dot representing a biological replicate that consists of 3 technical replicates. Bars on all graph indicate mean± SD. * = p < 0.05, ** = p < 0.01. Unpaired two-tailed t test with Welch’s correction (vs EV Control) was used for A. Multiple unpaired two-tailed t test with Welch’s correction and two-stage linear step-up procedure of Benjamini, Krieger and Yekutieli post-hoc testing was used for B (vs EV Control). Two-way ANOVA with two-stage linear step-up procedure of Benjamini, Krieger, and Yekutieli post-hoc testing was used for D and E (vs EV Control).

## Discussion

Two tissues that undergo similar stages of wound healing but with different outcomes are the oral mucosa and skin [1–3]. These tissue-specific healing response may be due to differential expressions of transcription factors [1, 6, 7]. We recently demonstrated that overexpression of BATF3, a transcription factor predicted to regulate the expression of genes involved in early palate wound healing, could improve wound-healing responses in HaCaT and induce differential expression of downstream genes [7]. CREB5 was another top transcriptional regulator in oral versus skin wounds and was constitutively expressed at higher levels in TIGK relative to HaCaT during *in vitro* healing [6]. Few studies have assessed *CREB5’s* role in wound healing or its influence on keratinocyte phenotype. Overexpression of CREB5 in HaCaT induced significan transcriptional changes and enhanced HaCaT migration.

Despite our prior study predicting that *SYNJ2* and *UBASH2B* would be positively regulated by *CREB5 in vivo* [7], they were significantly downregulated following CREB5 overexpression in HaCaT. This discordance may be due in part to the fact that our transcription factor prediction model is based on Microarray sequencing data from whole human skin, whereas our current experimental model uses only immortalized human keratinocyte cell lines *in vitro*. Future studies may use viral vectors to achieve more efficient transfection and possibly better-regulated CREB5 expression, thereby eliciting a more pronounced change in the HaCaT phenotype. It would be prudent to then examine phenotypic changes in HaCaT upon induction of multiple transcription factors predicted to be important for oral wound healing, as there may be combinatorial effects from multiple implicated transcription factors. Future studies involving transcription factor and gene regulatory network analyses should also utilize single-cell RNA sequencing data from human oral and skin tissues to obtain cell-type-specific data that may be more concordant with our *in vitro* models.

Consistent with our previous studies, our results suggest that transcription factors predicted to differentially regulate gene networks in early palate wounds compared with skin may contribute to the tissue-specific wound-healing outcomes [6, 7]. Moreover, inducing the expression of these transcription factors may enhance the wound-healing phenotype. Although many genes and pathways have been identified as vital to regenerative repair, we have yet to achieve tissue regeneration in human adult skin [1, 7, 13, 14]. Overall, our recent breadth of work demonstrates the value in identifying transcription factors and gene networks that distinguish oral from skin wound healing, as it may reveal targets to promote regenerative skin repair [6, 7]. This study also highlights two datasets that will be helpful for identifying candidate transcription factors.

