## Supplemental Table 1 for "A role for CREB5 in skin wound healing"

| Primer Sequences | | | |
|---|---|---|---|
| Gene | Forward Primer (5'-3') | Reverse Primer (5'-3") | Species |
| CREB5 | GGCTCTCTATCTTCTCTGC | GGAAGATCCTGGCATTGTAGGG | Human |
| CAMK2D | GCTTACTACAACCCTGCCAA | CTTCAAGCAGTCTACAGTCTCC | Human |
| TNFRSF10B | TGGACAGGACTATAGCACTCA | CTGTGTTTCTGGTCGTGGT | Human |
| SYNJ2 | ACTTTTGCAGACAGTCACTCG | ACCTTTCAGCCAGTCCTTG | Human |
| UBASH3B | AACATCTTCCCCCACATCAC | ACTTACATTTCCAGCGACTGAC | Human |
| ACTB | AGCACAGAGCCTCGCCTT | CATCATCCATGGTGAGCTGG | Human |

**Supplementary Table 1.** PCR primers used for RT-PCR of selected genes
